# Breastfeeding enhances intestinal gluconeogenesis in infants

**DOI:** 10.1101/2023.11.12.566732

**Authors:** Duan Ni, Jian Tan, Laurence Macia, Ralph Nanan

## Abstract

**Objectives:** Breastfeeding confers metabolic benefits to the infants, including reducing risks for metabolic syndromes such as obesity and diabetes later in life, but the underlying mechanisms is not yet fully understood. Hence, we aim to investigate the impacts of breastfeeding on the metabolic organs of infants.

**Methods:** Previous literatures directly studying the influences of breastfeeding on offspring’s metabolic organs were comprehensively reviewed. A microarray dataset of intestinal gene expression comparing infants fed on breastmilk versus formula milk was reanalyzed.

**Results:** Reanalysis of microarray data showed that breastfeeding significantly enhanced gluconeogenesis in infants’ intestine. This resembled observations in other mammalian species where breastfeeding also promoted gluconeogenesis.

**Conclusions:** Breastfeeding enhances intestinal gluconeogenesis in infants, which may underlie its metabolic advantages through finetuning the metabolic homeostasis. Such effect seems to be conserved across species, hinting its biological significance.

## Introduction

Breastfeeding (BF) delivers several advantages to infants. It programs their metabolic health, reducing the risk of obesity, diabetes and other metabolic syndromes later in life^1,2^. Studies investigating these associations have primarily been observational. A deeper mechanistic understanding of direct effects of BF on metabolic organs at a cellular level are lacking in humans. Here, we harnessed a published microarray dataset comparing intestinal gene expression between BF and formula feeding (FF) infants^3^. Comprehensive profiling their metabolic landscapes revealed enhanced intestinal gluconeogenesis (IGN) by BF. Our results align with studies in other mammalian species showing similar effects of BF on gluconeogenesis in metabolic organs. Since IGN contributes to modulating energy homeostasis, appetite and insulin sensitivity^4^, our data provides some potential mechanistic insights for the metabolic advantages reported in breastfed infants at a cellular level in humans.

## Methods

Details of gene expression and bioinformatic analysis were described in Supplementary Information.

## Results

We first collated evidence for cellular metabolism impacted by BF in non-human mammalian species (Figure 1A). In studies involving pigs, BF increased gluconeogenesis in offspring’s ileum and liver, evidenced by the upregulation of key genes in this pathway, like phosphoenolpyruvate carboxykinase 1 (PCK1). In breast-fed lambs, there was an increase in PCK2 expression in the rumen. Furthermore, the insulin receptor signaling pathway was found to be activated in the colon of breastfed mice.

**Figure 1.**
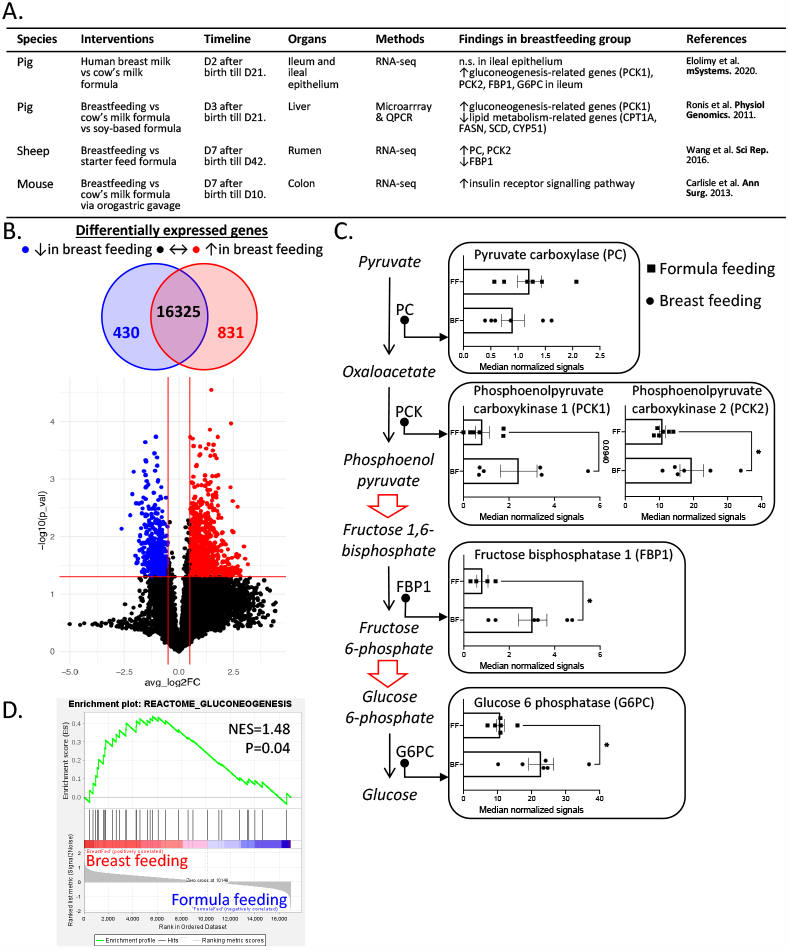
Breast feeding (BF) enhanced gluconeogenesis in infants from both animal models and humans. **A**. Overview of animal studies investigating the impacts of BF on the offspring’s metabolisms. **B**. Re-analysis of microarray data (GSE31075) revealed that BF upregulated 831 genes (red) and downregulated 430 genes (blue) in infants’ intestinal cells. **C**. Overview of the critical steps of gluconeogenesis pathway and the expression of critical genes involved in BF (round dots) versus formula feeding (FF, square dots) infants’ intestinal cells. **D**. Gene set enrichment analysis (GSEA) showed the enrichment of genes involved in gluconeogenesis pathways in BF (red) infants’ intestinal cells. (n.s.: no significant change; PCK1: phosphoenolpyruvate carboxykinase 1, PCK2: phosphoenolpyruvate carboxykinase 2, FBP1: fructose bisphosphatase 1, G6PC: glucose 6 phosphatase, CPT1A: carnitine palmitoyltransferase 1, FASN: fatty acid synthase, SCD: stearoyl-CoA desaturase, CYP51: cholesterogenic lanosterol 14alpha-demethylase, PC: pyruvate carboxylase, NES: normalized enrichment score)

Subsequently, leveraging a published microarray dataset, we comprehensively interrogated the impacts of BF versus FF on the intestinal metabolism of human infants. As shown in Supplementary Figure 1A, six BF and six FF infants, appropriately balanced, were analyzed. BF upregulated 831 and downregulated 430 genes respectively (Figure 1B). In-depth analysis unveiled several critical genes involved in gluconeogenesis, such as PCK2, fructose bisphosphatase 1 (FBP1) and glucose 6 phosphatase (G6PC), were increased by BF, implying enhanced IGN (Figure 1C). This was supported by Gene Set Enrichment Analysis, showing enrichment of gluconeogenesis genes in BF group (Figure 1D).

## Discussion

The intestine is an important metabolic organ, responsible for nutrient digestion and absorption. It is also a critical site for gluconeogenesis in addition to liver and kidney, maintaining metabolic homeostasis^4^. Here, we re-analyzed the intestinal microarray data from BF and FF infants and uncovered enhanced IGN by BF, possibly contributing to improved metabolic health. Importantly, the BF-induced IGN seems to be conserved across species, hinting its biological significance.

IGN is essential for maintaining overall glucose balance. It specifically decreases lipid accumulation in the liver, averting liver steatosis, and promotes thermogenesis in adipose tissues and the conversion of white fat to brown fat^4^. Additionally, glucose from IGN, when detected by the portal vein, can modulate food intake^4^. These processes may in part explain the metabolic advantages seen in BF infants, such as reduced risks of metabolic disorders later in life.

The exact mechanisms by which BF stimulates IGN remain elusive. While dietary protein is a known stimulant for IGN^4^, the slightly lower protein content in breast milk compared to infant formula suggests other factors to be at play. Alternatively, metabolites from the infants’ gut bacteria, like acetate and succinate, serve as important IGN fuels^5^. However, our microbiota analysis indicated so far only insignificant differences in their production pathways between BF and FF infants (Supplementary Figure 1), although further confirmations like metabolomics are needed. Conversely, breast milk itself is rich in maternal microbial metabolites^6^, which may explain the described changes.

Collectively, we provide compelling evidence for enhanced IGN triggered by BF in humans. These results shed light on the benefits of BF and justify further research to unravel the breast milk components responsible for IGN induction and to determine how long the increased IGN persists over time.

Duan Ni

Jian Tan, PhD

Laurence Macia, PhD

Ralph Nanan, Dr. med. Habil (Germany). FRACP

## Arthor Contributions

*Concept and design:* Duan Ni, and Ralph Nanan.

*Acquisition, analysis, and interpretation of data:* Duan Ni, Jian Tan, Laurence Macia, and Ralph Nanan.

*Drafting of the manuscript:* Duan Ni, and Ralph Nanan.

*Critical revision of the manuscript for important intellectual content:* All authors.

## Figure Legends

**Supplementary Figure 1.**
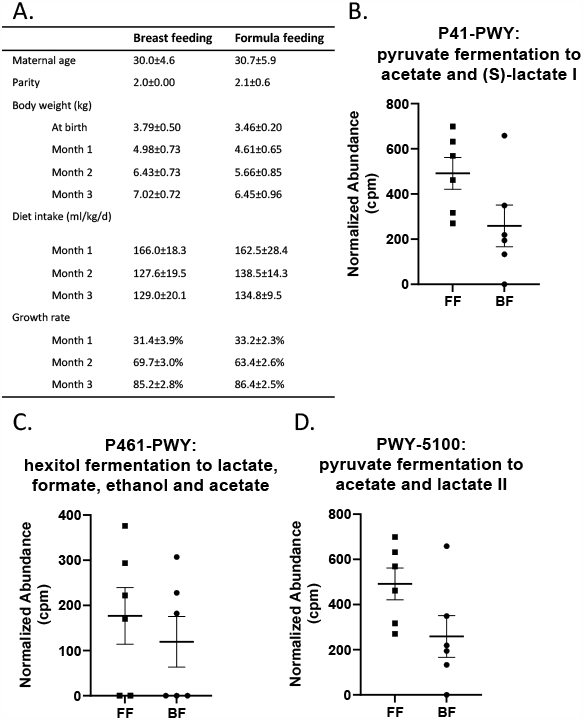
**A**. Overview of the breast feeding (BF) and formula feeding (FF) infants included in current study. (**B-D**) Scattering plots for the normalized abundance of the corresponding pathways predicted from microbiota analysis comparing BF (round dot) and FF (square dot) infants’ intestinal cells.

