## Supplementary Information for "Breastfeeding enhances intestinal gluconeogenesis in infants"

### Materials and Methods

Gene expression data was extracted from original papers shown in Figure 1A. Normalized microarray data was downloaded from [1] (GEO31075) and analyzed with R or PRISM GraphPad. Gene set enrichment analysis (GSEA) was run following the software's tutorial.

For metagenomic analysis, raw sequence minus human-identical sequences was downloaded from [1] (ERP001038). After quality filtering, sequence was clustered using CDHIT-454 and taxonomic assignment on resulting reads were performed using MetaPhlAn2. The HUMAnN2 pipeline was utilized for functional profiling to identify gene families and MetaCyc metabolic pathways.
